# High-resolution mapping of *Rym14^Hb^*, a wild relative resistance gene to barley yellow mosaic disease

**DOI:** 10.1101/2020.08.06.239145

**Authors:** Hélène Pidon, Neele Wendler, Antje Habekuβ, Anja Maasberg, Brigitte Ruge-Wehling, Dragan Perovic, Frank Ordon, Nils Stein

## Abstract

Barley yellow mosaic disease is caused by Barley yellow mosaic virus and Barley mild mosaic virus, and leads to severe yield losses in barley (*Hordeum vulgare*) in Central Europe and East-Asia. Several resistance loci are used in barley breeding. However, cases of resistance-breaking viral strains are known, raising concerns about the durability of those genes. *Rym14*^*Hb*^ is a dominant major resistance gene on chromosome 6HS, originating from barley’s secondary genepool wild relative *Hordeum bulbosum*. As such, the resistance mechanism may represent a case of non-host resistance, which could enhance its durability. A susceptible barley variety and a resistant *H. bulbosum* introgression line were crossed to produce a large F_2_ mapping population (n=7,500), to compensate for a ten-fold reduction in recombination rate compared to intraspecific barley crosses. After high-throughput genotyping, the *Rym14*^*Hb*^ locus was assigned to a 2Mbp telomeric interval on chromosome 6HS. The co-segregating markers developed in this study can be used for marker-assisted introgression of this locus into barley elite germplasm with a minimum of linkage drag.

## Introduction

Viruses are an increasing threat to crops worldwide. The soil-borne barley yellow mosaic disease, caused by a complex of two *Bymoviruses* (*Barley yellow mosaic virus* (BaYMV) and *Barley mild mosaic virus* (BaMMV)) is one of the most important diseases of winter barley. Widespread in central Europe and East Asia, it causes severe yield losses up to even total crop failure (Plumb et al. 1986; Jianping 2005; Kühne 2009). As chemical control of those viruses, transmitted by the plasmodiophorid *Polymyxa graminis* (Kanyuka et al. 2003), is not possible, only the use of resistant varieties can preserve yield in infected fields.

To date, 20 barley resistance genes have been identified, almost exclusively conferring recessive resistance (Jiang et al. 2020). Two of these loci have been cloned: the *EUKARYOTIC TRANSLATION INITIATION FACTOR 4E* gene (*eIF4E*), (Stein et al. 2005) of which several allelic forms providing resistance are described, including *rym4* and *rym5*, (Hofinger et al. 2011; Perovic et al. 2014; Yang et al. 2017; Shi et al. 2019), and the *PROTEIN DISULFIDE ISOMERASE LIKE 5-1* (*PDI5-1*) gene which is also represented by a handful of alleles providing resistance, including *rym1* and *rym11* (Yang et al. 2017). The *rym4* allele provides resistance to BaMMV and to the common BaYMV pathotype BaYMV-1, but not to pathotype BaYMV-2, which emerged in Europe at the end of the 1980s (Adams et al. 1987; Huth 1989; Adams 1991; Graner and Bauer 1993; Steyer et al. 1995). The spectrum of *rym5* covers also *BaYMV-*2, however, resistance-breaking isolates of BaMMV and BaYMV have emerged (Kanyuka et al. 2004; Habekuß et al. 2008; Li et al. 2016). Facing the prospect of boom-and-bust cycles for known resistance genes (Brown and Tellier 2011), it is critical to continue searching for alternative resistance loci to underpin resistance breeding and to allow pyramiding of disease resistance loci. In particular, sources of non-host resistance, e.g. resistance exhibited from a plant species against all isolates of a pathogen which is not coevolutionary adapted, are particularly promising as they are thought to cover a larger resistance spectrum and to be more durable (Ayliffe and Sørensen 2019). Bulbous barley (*Hordeum bulbosum* L.), a wild relative and representative of the secondary gene pool of cultivated barley (*Hordeum vulgare* L.), has been described as source of resistance to numerous barley pathogens, including barley leaf rust (Johnston et al. 2013; Yu et al. 2018) and barley powdery mildew (Xu and Kasha 1992; Pickering et al. 1995; Shtaya et al. 2007). So far, all *H. bulbosum* accessions investigated exhibited resistance to BaMMV and BaYMV (Ruge et al. 2003), suggesting that the species is probably a non-host to those viruses. Two major dominant resistance genes from *H. bulbosum* to both BaMMV and BaYMV have been described: *Rym14*^*Hb*^ (Ruge et al. 2003) and *Rym16*^*Hb*^ (Ruge-Wehling et al. 2006). *Rym14*^*Hb*^ was introgressed to barley by translocation of a *H. bulbosum* segment to barley chromosome 6HS (Ruge et al. 2003). In the past, a lack of suitable markers, alongside severely reduced recombination in the target region between the barley and *H. bulbosum* fragments, rendered precise mapping of *Rym14*^*Hb*^ elusive. Thanks to the development of genetic and genomic resources for *H. bulbosum* (Wendler et al. 2014, 2015), it is now possible to fine-map loci from this species in a *H. vulgare* background.

We aimed to map *Rym14*^*Hb*^ at high resolution, and to provide markers for its introgression into elite barley, ideally without linkage drag, using large populations and high-throughput genotyping to overcome the lack of recombination.

## Materials and methods

### Plant material

A first round of low-resolution genetic mapping was performed using four F_6_ families derived from F_5_ plants heterozygous at the *Rym14*^*Hb*^ locus from the BAZ-4006 family of the population ‘Borwina’ x ‘A42’ described in Ruge et al. (2003).

To achieve a population size suitable for fine mapping, an additional eight F_2_ families were generated by crossing an *Rym14*^*Hb*^/*Rym14*^*Hb*^ F_6_ plant (derived from F_5_ 4006/337) to either (i) var. ‘KWS Orbit’ or (ii) var. ‘KWS Higgins’, both missing the *Rym14*^*Hb*^ resistance locus (-/-). In the purpose of instant pyramiding of disease resistance loci both cultivars carry *rym4*-based resistance (*rym4*/*rym4*) to BaMMV and BaYMV.

### DNA extraction

Genomic DNA of plants from the low-resolution mapping population was isolated as described by Stein et al. (2001). Genomic DNA of plants from the fine-mapping population was extracted according to the guanidine isothiocyanate-based protocol described by Milner et al. (2019).

### Genotyping-by-sequencing and data analysis

GBS libraries for the low-resolution mapping were prepared from genomic DNA digested with *Pst*I and *Msp*I (New England Biolabs) as described by Wendler et al. (2015). Between 93 and 153 barcoded samples were pooled in an equimolar manner per lane and sequenced on the Illumina HiSeq 2500 for 107 cycles, single-end reads, using a custom sequencing primer.

The GBS reads were processed, aligned, and used to generate variant calls as described by Milner et al. (2019). Alignment was performed against the TRITEX genome assembly of barley cultivar ‘Morex’ (Monat et al. 2019). Individual variant calls were accepted wherever the read depth exceeded four. Variant sites were retained if they presented a minimum mapping quality score (based on read depth ratios calculated from the total read depth and depth of the alternative allele) of 20, a maximum fraction of 40% of missing data, a fraction of heterozygous calls between 30 and 70%, and between 10 and 40% of each homozygous call. Individuals with more than 40% missing data were excluded.

### Marker development

Exome capture data of the introgression line ‘4006/163’, described in Wendler et al. (2014) (accession number ERP004445), were mapped to the TRITEX genome assembly of barley cultivar Morex (Monat et al. 2019) together with the exome capture data of the *H. bulbosum* genotype ‘A42’ and of eight barley varieties: ‘Bonus’, ‘Borwina’, ‘Bowman’, ‘Foma’, ‘Gull’, ‘Morex’, ‘Steptoe’, and ‘Vogelsanger Gold’, described in Mascher et al. (2013b) (accession number PRJEB1810). Read mapping and variant calling were performed as described by Milner et al. (2019). The SNP matrix was filtered for the following criteria: heterozygous and homozygous calls had to be covered by a minimum depth of three and five reads, respectively, and have a minimum quality score of 20. SNP sites were retained if they had less than 20% missing data and less than 20% heterozygous calls. SNPs that were carrying the reference call in all eight barleys and the alternate call in ‘A42’ and ‘4006/163’ were selected as candidates to design KASP markers, either using KASP-by-design (LGC Genomics, Berlin, Germany) or 3CR Bioscience (Essex, UK) free assay design service. Those markers are latter designated as KASP and PACE markers, respectively. Since no suitable SNPs were identified in the first 500 kbp of chromosome 6HS on the ‘Morex’ reference genome, the exome capture data were additionally mapped to the genome assembly of cultivar ‘Barke’ (Jayakodi et al. under revision). The SNP at coordinate 241,723 bp on chromosome 6H of the ‘Barke’ genome assembly was retrieved and used to design the telomeric marker Rym14_Bar241723. Furthermore, in order to control the genetic state at the segregating *rym4* resistance locus, the diagnostic SNP for the resistance conferring allele (Stein et al. 2005) was also used to design a KASP marker. Further information on KASP and PACE markers is provided in supplementary tables 1 and 2, respectively.

**Table 1.** Genes annotated with high confidence in *Rym14*^*Hb*^ interval on the Morex genome (Monat et al. 2019).

| name | start | stop | gene type |
| --- | --- | --- | --- |
| HORVU.MOREX.r2.6HG0447840 | 195540 | 196334 | Thionin |
| HORVU.MOREX.r2.6HG0447850 | 220610 | 221213 | Thionin |
| HORVU.MOREX.r2.6HG0447860 | 256998 | 259999 | Thionin |
| HORVU.MOREX.r2.6HG0447880 | 373994 | 438209 | Thionin |
| HORVU.MOREX.r2.6HG0447890 | 460556 | 461157 | Thionin |
| HORVU.MOREX.r2.6HG0447900 | 461856 | 462457 | Thionin |
| HORVU.MOREX.r2.6HG0447910 | 497194 | 497795 | Thionin |
| HORVU.MOREX.r2.6HG0447920 | 597800 | 598403 | Thionin |
| HORVU.MOREX.r2.6HG0447930 | 625302 | 625905 | Thionin |
| HORVU.MOREX.r2.6HG0447940 | 691184 | 707575 | Thionin |
| HORVU.MOREX.r2.6HG0447950 | 749829 | 776991 | Thionin |
| HORVU.MOREX.r2.6HG0447960 | 792195 | 827832 | Thionin |
| HORVU.MOREX.r2.6HG0447980 | 958137 | 958736 | Thionin |
| HORVU.MOREX.r2.6HG0447990 | 1004017 | 1004618 | Thionin |
| HORVU.MOREX.r2.6HG0448010 | 1259976 | 1260591 | TIR-NBS-LRR class disease resistance protein |
| HORVU.MOREX.r2.6HG0448020 | 1300107 | 1300565 | Dimeric alpha-amylase inhibitor |
| HORVU.MOREX.r2.6HG0448100 | 1493250 | 1493945 | Dirigent protein |
| HORVU.MOREX.r2.6HG0448110 | 1574160 | 1575749 | Cytochrome P450 family protein, expressed |
| HORVU.MOREX.r2.6HG0448120 | 1578752 | 1580023 | Aspartic proteinase nepenthesin-1 |
| HORVU.MOREX.r2.6HG0448130 | 1598418 | 1600649 | Subtilisin-like protease |
| HORVU.MOREX.r2.6HG0448140 | 1605306 | 1610732 | Fatty acyl-CoA reductase |
| HORVU.MOREX.r2.6HG0448160 | 1753412 | 1756451 | Glycerol-3-phosphate acyltransferase 3, putative |
| HORVU.MOREX.r2.6HG0448200 | 1792383 | 1794963 | Transposon protein, putative, CACTA, En/Spm sub-class |
| HORVU.MOREX.r2.6HG0448210 | 1796825 | 1804280 | O-acyltransferase WSD1 |
| HORVU.MOREX.r2.6HG0448220 | 1840897 | 1842376 | GDSL esterase/lipase |
| HORVU.MOREX.r2.6HG0448230 | 1853483 | 1854626 | Short-chain dehydrogenase/reductase |
| HORVU.MOREX.r2.6HG0448250 | 1945996 | 1952442 | Protein kinase family protein |
| HORVU.MOREX.r2.6HG0448260 | 1954346 | 1955384 | zinc finger MYM-type-like protein |
| HORVU.MOREX.r2.6HG0448290 | 2061596 | 2062919 | Cysteine protease-like protein |
| HORVU.MOREX.r2.6HG0448300 | 2066856 | 2067293 | Proteinase inhibitor type-2 |

### Genotyping

Genotyping assays with KASP markers were carried out in a final volume of 5 μl consisting of 0.7 μl genomic DNA (50-100 ng/µL), 2.5 μl of KASP V4.0 2X Master Mix High Rox (LGC Genomics, Berlin), 0.07 μl KASP assay mix (KASP-by-design, LGC Genomics, Berlin) containing the primers, and 2.5 μl of sterile water. PCR amplifications were performed using the Hydrocycler 16 (LGC Genomics, Berlin) with cycling conditions as follows: 94 °C for 15 min, followed by a touchdown profile of 10 cycles at 94 °C for 20 s and 61 °C for 1 min with a 0.6 °C reduction per cycle, followed by 26 cycles at 94 °C for 20 s and 55 °C for 1 min. Genotyping assays with PACE markers were carried out in a final volume of 5 μl consisting of 0.7 μl genomic DNA (50-100 ng/µL), 2.5 μl of PACE Master Mix High Rox (3cr Bioscience, Essex, United Kingdom), 0.07 μl primer mix containing the primers (12 µM of each allele specific primers and 30 µM of the common reverse primer), and 2.5 μl of sterile water. PCR amplifications were performed using the Hydrocycler 16 (LGC Genomics, Berlin) with cycling conditions as follows: 94 °C for 15 min, followed by a touchdown profile of 10 cycles at 94 °C for 20 s and 65 °C for 1 min with a 0.8 °C reduction per cycle, followed by 30 cycles at 94 °C for 20 s and 57 °C for 1 min.

For both marker types, the genotyping results were read out using the ABI 7900HT (Applied Biosystems) using an allelic discrimination file. Readings were made before and after PCR, and the data were analyzed using SDS 2.4 Software (Applied Biosystems).

### Phenotyping

Resistance to BaMMV was tested under greenhouse conditions as described by Habekuß et al. (2008). After sowing, the plants were grown in a greenhouse (16-h day/8-h night, 12 °C). The susceptible barley variety ‘Maris Otter’ was systematically included to monitor success of infection. At the 3-leaf stage (around 2 weeks after sowing), the plants were mechanically inoculated twice at an interval of 5–7 days with the isolate BaMMV-ASL1 (Timpe and Kühne 1994) using the leaf-sap of BaMMV-infected leaves of susceptible cv. ‘Maris Otter’, mixed in K_2_HPO_4_ buffer (1:10; 0.1 M; pH 9.1) containing silicon carbide (caborundum, mesh 400, 0.5 g/25 ml sap). Five weeks after the first inoculation, the number of infected plants with mosaic symptoms were scored, and DAS-ELISA with BaMMV-specific antibodies was carried out in parallel according to published protocols (Clark and Adams 1977). Virus particles were estimated via extinction at 405 nm using a Dynatech MR 5000 microtiter-plate reader. Plants with an extinction E_405_>0.1 were qualitatively scored as susceptible.

## Results

### Low-resolution mapping

A population of 427 F_6_ from the cross ‘Borwina’ x ‘A42’ was genotyped by GBS and phenotyped for resistance to BaMMV. Data for 389 plants and 77 SNPs passed the quality filters (supplementary table 3). On chromosome 6H, 73 plants were homozygous for the ‘Borwina’ allele, 92 were homozygous for the ‘A42’ allele, 220 were heterozygous and four recombined. The infection rate was low with only 10% of plants infected, compared to an expected 25% when resistance is controlled by a single dominant gene. However, among the 39 plants phenotyped as susceptible to BaMMV, 38 were homozygous for the ‘Borwina’ allele and one recombined on chromosome 6H, indicating a strong association of phenotype and genotype.

To further confirm this association, 26 lines were phenotyped on progenies of 12 to 20 plants (Figure 1, supplementary table 4). These included (i) 17 lines with the susceptible genotype on chromosome 6H but scored as resistant, (ii) five heterozygous lines, and (iii) the four recombinant lines. Progenies of lines presenting the susceptible genotype displayed infection rates between 50 and 95%, while those of heterozygous lines displayed rates between 5 and 17%.

**Fig. 1.**
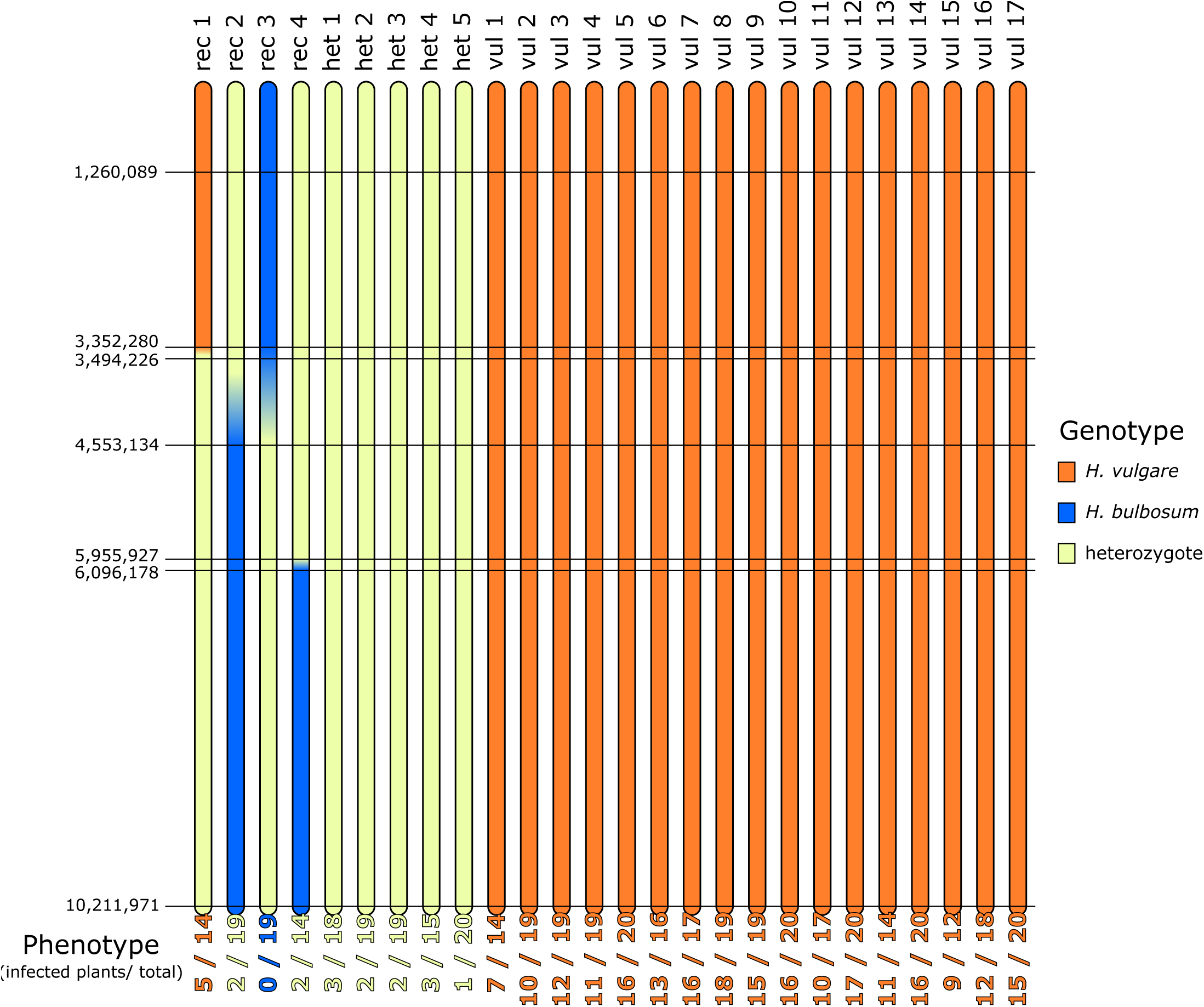
Graphical genotype and phenotype of the 26 F_6_ lines phenotyped on progenies. *H. vulgare, H. bulbosum*, and heterozygous phenotyped are represented as orange, blue, and yellow bars, respectively. Coordinates on Morex reference genome (Monat et al. 2019) of strategic markers are displayed. Phenotypes are indicated as the number of infected plants out of the total of F_7_ progenies phenotyped, colored according to the F_6_ phenotype interpreted, following the same color code as for genotypes.

These results support the low penetrance of the infection in this experiment, with only half of the expected susceptible plants successfully infected, as well as the association of the chromosome 6H locus with resistance to BaMMV. Moreover, the phenotypes of the four recombinant progenies defined *Rym14*^*Hb*^ interval between the telomere of chromosome 6HS and the marker position at base pair 4,553,134.

### Fine mapping

The population of 7,500 F_2_ was genotyped at the *Rym14*^*Hb*^ locus with four KASP markers (Rym14_Bar241723, Rym14_2370223, Rym14_3087282, and Rym14_5003183, supplementary table 1). Resistance due to segregation of the recessive resistance gene *rym4* on chromosome 3HL was controlled for with the rym4_SNP KASP marker (supplementary table 1). We identified 28 recombination events, corresponding to a genetic distance of ∼0.2 cM, between the markers Rym14_Bar241723 and Rym14_5003183. These results confirmed the strongly reduced recombination rate between the *H. bulbosum* and the *H. vulgare* fragments on chromosome 6HS. In cultivated barley, the syntenic 5 Mbp *Rym14*^*Hb*^ interval on chromosome 6HS corresponds to a genetic distance of 4 cM (Mascher et al. 2013a), implying a 20-fold reduction in recombination frequency between the *H. bulbosum* and the *H. vulgare* fragment.

All recombinants were genotyped with seven PACE markers (supplementary tables 2 and 4). Among the recombinants, ten plants were homozygous for the *rym4* allele, nine were heterozygous and the remaining nine were homozygous wildtype at the *rym4* locus (supplementary table 5). As plants homozygous for the *rym4* allele would be resistant to BaMMV, irrespective to their genotype at *Rym14*^*Hb*^, only F_3_ families derived from the 18 *Rym14*-recombinants heterozygous or homozygous for the susceptible allele at *rym4* were phenotyped using 30 and 20 F_3_ siblings, respectively. All phenotyped plants were genotyped at Rym14_Bar241723, Rym14_2370223, Rym14_5003183 and *rym4* (supplementary table 6). The infection rate during this round of phenotyping was much higher than during the preceding low-resolution mapping, with less than 2 % of the susceptibility control showing no viral content. Five out of 86 F_3_ siblings expected to be susceptible based on their genotype were not infected by virus, hence producing false-negative phenotypic results.

Based on this analysis, the *Rym14*^*Hb*^ target region was reduced to a 2 Mbp interval on the Morex reference genome, between the telomere of chromosome 6HS and Rym14_2066975 (figure 2).

**Fig. 2.**
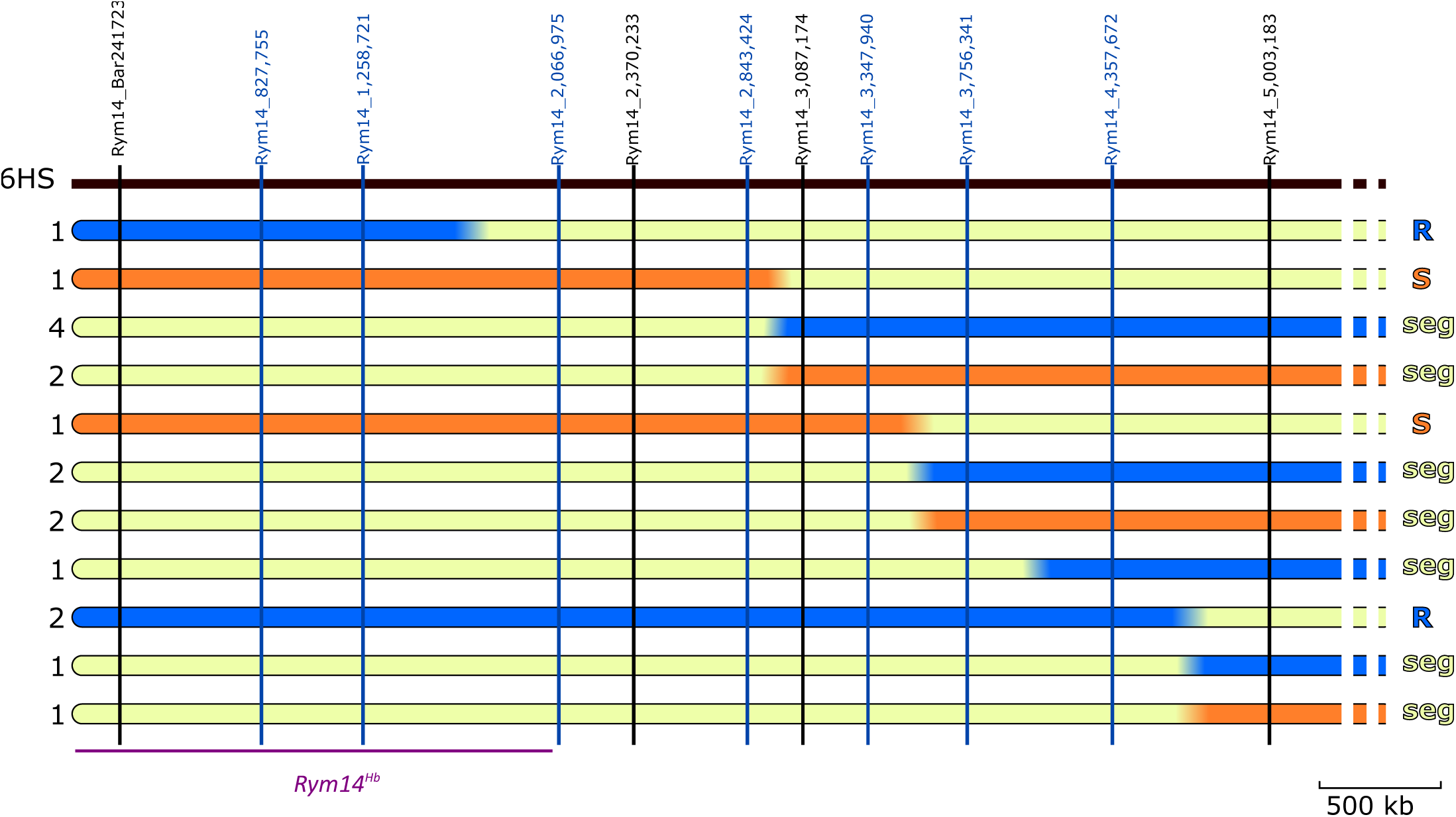
Physical map of the *Rym14*^*Hb*^ locus. KASP and PACE markers are represented as black and blue vertical lines, respectively. Barley chromosome 6HS is depicted as a black horizontal line and genotypes of recombinant F_2_ plants are indicated by horizontal bars: blue=*H. bulbosum* homozygous; orange=*H. vulgare* homozygous; yellow=heterogygous. The number of recombinant lines corresponding to each genotype pattern is indicated on the left while the phenotypes of their progeny are shown on the right (R: resistant, S: susceptible, seg: segregation of resistance).

### Candidate genes

In the absence of a genomic sequence for a *Rym14*^*Hb*^ plant, we cannot precisely define the genes present in the *Rym14*^*Hb*^ interval. However, as synteny between the two *Hordeum* species is high (Wendler et al. 2017), it is still relevant to assess the genes annotated in the homolog interval of the *H. vulgare* reference genome as a proxy for suggesting *Rym14*^*Hb*^ candidate genes. In the respective interval of the Morex V2 reference sequence 30 high-confidence (HC) (Table 1) and 17 low-confidence genes (Monat et al. 2019) are annotated. All HC gene models were checked for homology with other genes by a BLASTx (v2.9.0, default parameters) homology searches against the non-redundant protein sequence database (Camacho et al. 2009) and for presence of conserved domains in NCBI conserved domains (Lu et al. 2019). Among the HC genes, HORVU.MOREX.r2.6HG0448010 is annotated as a TIR-NBS-LRR gene, however, it does not contain any of the major NLR domains (coiled-coil, NB-ARC and LRR), and is therefore interpreted as a pseudogene. HORVU.MOREX.r2.6HG0448100, annotated as a dirigent protein, is a jacalin-related lectin, while HORVU.MOREX.r2.6HG0448250, annotated as part of the protein kinase protein family, displays the highest homology with a wall-associated receptor kinase, and HORVU.MOREX.r2.6HG0448290 codes for a papain-like cysteine protease (PLCP). Interestingly, the interval also contains no less than 14 HC genes annotated as thionins, sharing with each other at least 88% of their coding sequence. In addition to these annotated genes in the Morex genome, additional candidate genes could be unique to the resistant genotypes.

## Discussion

Resistance genes deployed in breeding and in the field are often overcome by new pathogen variants after only a few years (Brown and Tellier 2011). Pyramiding several resistance genes has proven to increase the resistance durability, however, this strategy requires the availability of several independent resistance loci (Werner et al. 2005; Riedel et al. 2011; Kim et al. 2011). In light of these facts, non-adapted resistance genes from wild crop relatives are precious, since they are assumed to confer more durable resistance than genes originating from within the diversity of the cultivated species, owing to co-evolution between the cultivated host and pathogen genotypes (Fonseca and Mysore 2019). Until recently, the fine mapping of genes from crop wild relatives species was impractical, owing to strong suppression of recombination with the cultivated species (Ruge et al. 2003; Kakeda et al. 2008; Wijnker and de Jong 2008; Prohens et al. 2017). The results of this study demonstrate that high-throughput genotyping coupled with large mapping populations can overcome this limitation, by constraining the interval of the *Rym4*^*Hb*^ viral resistance gene to the telomeric 2 Mbp of chromosome 6HS, and providing markers suitable for marker-assisted-selection.

While genes coding for nucleotide-binding and leucine-rich repeat domain proteins (NLR) are the usual suspects for dominant resistance to pathogens, including viruses (de Ronde et al. 2014; Boualem et al. 2016), only a pseudogene presenting similarities with this gene family is annotated in the *Rym14*^*Hb*^ interval on the barley reference genome. However, it is not rare that susceptible genotypes do not possess a functional copy of the resistance gene. NLRs are overrepresented in regions displaying presence/absence variation (Xu et al. 2012; Bush et al. 2013). Therefore, some NLR resistance genes, like *RPM1* and *RPS5*, are only present in the resistant genotype (Grant et al. 1998; Henk et al. 1999). In the case of wheat leaf rust resistance gene *Lr21*, it was shown that the gene is a chimera of two nonfunctional alleles that probably evolved via a recombination event (Huang et al. 2009).

Among the other annotated genes at the *Rym14*^*Hb*^ locus, two are very good candidates. Wall-associated protein kinase-like HORVU.MOREX.r2.6HG0448250 are described resistance genes in plant-bacteria and plant-fungus pathosystems (Li et al. 2009, 2020; Dmochowska-Boguta et al. 2020). Their role in plant-virus pathosystems is less clear but it has been suggested that a cell wall-associated protein kinase was involved in the repression of plasmodesmal transport of the Tobacco mosaic virus by phosphorylating its movement protein (Citovsky et al. 1993; Waigmann et al. 2000). A second promising candidate is HORVU.MOREX.r2.6HG0448100. It codes for a jacalin-related lectin and is thus part of the family that includes the *Arabidopsis thaliana* genes *RTM1* and *JAX1* that provide dominant major resistance against potyviruses and potexviruses, respectively (Chisholm et al. 2000; Yamaji et al. 2012).

However, other genes in the *Rym14*^*Hb*^ interval, even if less likely candidates, might also play a role in resistance. For example, HORVU.MOREX.r2.6HG0448290 codes for a PLCP. PLCPs are known to play a major role in programmed cell death triggered by NLR genes. Interestingly, CYP1, a tomato PLCP, is targeted by the Tomato yellow leaf curl virus V2 protein, suggesting that V2 could downregulate CYP1 to counteract host defenses (Bar-Ziv et al. 2012). *Rcr3*, a tomato papain-like cysteine protease gene, is required for the function of the resistance gene *Cf-2* to *Cladosporium fulvum* (Krüger et al. 2002), while *NbCathB*, from *Nicotina benthamiana*, is requested for the HR triggered by the non-host pathogens *Erwinia amylovora* and *Pseudomonas syringae* (Gilroy et al. 2007). The high level of thionin duplication at this locus also raised our attention. Thionins are part of common anti-bacterial and anti-fungal peptides (Bohlmann and Broekaert 1994), conferring enhanced resistance to several pathogens. Thionins were also found to exhibit increased expression in resistant compared to susceptible pepper genotypes during infection by the Chili leaf curl virus (Kushwaha et al. 2015), suggesting a possible role in basal defense. Additionally, the cytochrome P450 superfamily has been associated with resistance to the Soybean mosaic virus (Cheng et al. 2010; Yang et al. 2011). Some subtilisin proteases are induced by pathogens and involved in programmed cell death (Figueiredo et al. 2014), and GDSL lipases were found to be either negative or positive regulators of plant defense mechanisms (Hong et al. 2008; Kwon et al. 2009).

The feasibility of further reducing the target interval by recombination through additional fine mapping is low and would require the screening of tens of thousands of additional F_2_ plants for the chance of finding one additional recombinant in the smallest target region. Therefore, a candidate gene approach may be a more fruitful strategy for continued progress. Despite the presence of promising candidate genes like HORVU.MOREX.r2.6HG0448250 and HORVU.MOREX.r2.6HG0448100 in the haplotype of the susceptible cultivar Morex, the resistance conferring gene may be present only in the haplotype of the resistant *H. bulbosum*. Therefore, deciphering the resistant haplotype, most likely though a high-quality chromosome-scale genome assembly of the interval in *H. bulbosum*, is an essential prerequisite to the prioritization of candidate genes for further functional testing.

The markers identified in this study are tightly linked to *Rym14*^*Hb*^ and therefore are of prime importance to barley breeding. These markers will allow the reliable introgression of this resistance into barley elite lines with a minimum of linkage drag compared to the previously established markers (Ruge et al. 2003). This is essential for introducing this gene into new cultivars, As the prevalence of resistance-breaking isolates of *rym4* and *rym5* will increase in the barley growing area in Europe and Asia (Kühne 2009), introgression of *Rym14*^*Hb*^ into new elite varieties together with other resistance loci represents a critical opportunity to improve the durability and spectrum of barley resistance to BaMMV and BaYMV.

## Declarations

### Conflicts of interest/Competing interest

NW and AM are employed at KWS SAAT SE & Co and KWS LOCHOW, respectively. The other authors declare no conflict of interest.

### Ethics approval

Not applicable

### Consent to participate

Not applicable

### Consent for publication

Not applicable

### Availability of data and material

The GBS dataset generated and analyzed in this study is deposited at EMBL-ENA under the project ID PRJEB39211 (not accessible during peer-review).

### Code availability

Not applicable

## Acknowledgments

We gratefully acknowledge the excellent technical support by Manuela Kretschmann in DNA extraction and KASP genotyping, Dörte Grau in BaMMV resistance phenotyping and Susanne König in GBS library preparation. We thank Axel Himmelbach for his valuable support in next generation sequencing, Klaus Oldach, Viktor Korzun and Jörg Grosser for their valuable inputs and Timothy Rabanus-Wallace for language editing.

## Key message

We mapped the *Rym14*^*Hb*^ resistance locus to barley yellow mosaic disease in a 2Mbp interval. The co-segregating markers will be instrumental for marker assisted selection in barley breeding.

## Supplementary material

**Table S1** KASP markers developed for *Rym14*^*Hb*^ fine mapping. The indicated coordinates of the genotyped SNP is respective to Morex V2 genome (Monat et al. 2019), except for Rym14_Bar241723 which it is based on Barke assembly (Jayakodi et al. under revision). The target SNP is identified in the sequence by square brackets.

**Table S2** PACE markers developed for *Rym14*^*Hb*^ fine mapping. The indicated coordinates of the genotyped SNP is respective to Morex V2 genome (Monat et al. 2019). The target SNP is identified in the sequence by square brackets.

**Table S3** Phenotype and filtered GBS genotype of 389 F_6_ plants from the cross Borwina x A42. Phenotype is either resistant (R) or susceptible (S). For each SNP, the genotype is indicated as homozygous *H. bulbosum* (B), homozygous *H. vulgare* (V) or heterozygous (H) and missing (-).

**Table S4** Phenotype on F_2_, phenotype on progenies and filtered GBS genotype of 26 lines from the cross Borwina x A42. Phenotype is either resistant (R) or susceptible (S). The number of susceptible plants out of the total number phenotyped for each progeny is specified. For each SNP, the genotype is indicated as homozygous *H. bulbosum* (B), homozygous *H. vulgare* (V) or heterozygous (H) and missing (-).

**Table S5** Genotyping of the 28 F_2_ recombinants with PACE and KASP markers. For each *Rym14*^*Hb*^ marker, the genotype is indicated as homozygous *H. bulbosum* (B), homozygous *H. vulgare* (V) or heterozygous (H). Genotype at *rym4* locus is classified as homozygous *rym4* (rym4_R), homozygous for the susceptible allele (rym4_S) and heterozygous (rym4_H). Additionally, the number of susceptible and resistant plants in the phenotyped progenies is specified.

**Table S6** Phenotype and genotyped of the F_3_ progenies recombining at the *Rym14*^*Hb*^ locus. The phenotype is given as the DAS-ELISA extinction at 405 nm. Plants with absorbance > 0.1 were scored qualitatively as being susceptible. For each *Rym14*^*Hb*^ marker, the genotype is indicated as homozygous *H. bulbosum* (B), homozygous *H. vulgare* (V) or heterozygous (H). Genotype at *rym4* locus is classified as homozygous *rym4* (rym4_R), homozygous for the susceptible allele (rym4_S) and heterozygous (rym4_H).

## Notes

**Funding** This work was supported as part of the collaborative projects “TransBulb” (grant 0315966 from the German Federal Ministry of Education and Research (BMBF)) and “BulbOmics” (grant 2818201615 from the German Federal Ministry of Food and Agriculture (BMEL)).

